# Dynamic modulation of IRE1α-XBP1 signaling by adenovirus

**DOI:** 10.1101/2024.11.30.626188

**Authors:** Yumi Jang, Fred Bunz

## Abstract

The abundant production of foreign proteins and nucleic acids during viral infection elicits a variety of stress responses in host cells. Viral proteins that accumulate in the endoplasmic reticulum (ER) can trigger the unfolded protein response (UPR), a coordinated signaling program that culminates in the expression of downstream genes that collectively restore protein homeostasis. The model pathogen adenovirus serotype 5 (HAdV5) activates the UPR via the signaling axis formed by inositol-requiring enzyme type 1 (IRE1α) and the X-box binding protein 1 (XBP1), a transcription factor required for immune function. Recent studies have suggested that IRE1α-XBP1 activity supports adenovirus replication. Here, we show that HAdV5 exerted opposing effects on IRE1α and XBP1. IRE1α was activated in response to HAdV5 but the production of the XBP1 isoform, XBP1s, was post-transcriptionally blocked. The tumor suppressor p53, which is eliminated by HAdV5 after infection, inhibited IRE1α activation. The de-repression of IRE1α following the degradation of p53 conceivably reflects a novel antiviral mechanism, which HAdV5 ultimately evades by suppressing XBP1s. Our findings highlight the defective antiviral defenses in cancer cells and further illustrate the opposing mechanisms used by adenoviruses and their host cells to exert control over the UPR, a critical determinant of cell fate.

## Introduction

Many disease states, including infections and cancer, are characterized by an intracellular overabundance of unfolded and misfolded proteins. Under normal physiologic conditions, newly synthesized proteins destined for secretion or incorporation into membranes are properly folded by molecular chaperones in the endoplasmic reticulum (ER).^1,2^ The protein processing capacity of the ER can be overwhelmed by pathologic levels of transcription and/or translation. Protein overload and other causes of ER stress trigger the activation of UPR signaling pathways that orchestrate a return to homeostasis. The UPR can expand the protein-folding capacity of the ER, increase protein turnover and protein secretion, and inhibit protein translation. Should these measures fail to restore homeostasis, downstream mediators of the UPR can trigger programmed cell death.

The IRE1α-XBP1 pathway is the most evolutionarily ancient of the three distinct signaling axes that mediate the UPR, and is conserved in all eukaryotes.^3^ Signals triggered by unfolded proteins in the ER lumen are initially generated at the ER membrane by IRE1α, a transmembrane protein with protein kinase and RNase moieties.^4^ When ER stress is below an activating threshold, the luminal domain of IRE1α is stably associated with a chaperone called BiP. An increase in the abundance of unfolded proteins in the ER lumen causes the translocation of BiP from IRE1α to client proteins and thereby permits inactive IRE1α monomers to self-associate into active oligomers.^5^

The cytoplasmic domain of activated IRE1α catalyzes the noncanonical splicing of the ER-associated RNA transcript expressed from the *XBP1* locus (Fig 1A). The unspliced *XBP1* transcript encodes XBP1u, an ephemeral protein of uncertain function.^6^ By excising a 26 bp intron, IRE1α produces a frameshift in the *XBP1* coding sequence. This spliced transcript directs the expression of a stable transcription factor called XBP1s. Transported into the nucleus, XBP1s alleviates ER stress and functions broadly in processes including lipid metabolism, glucose homeostasis and inflammation through the transactivation of a diverse set of target genes.^7^

**Figure 1.**
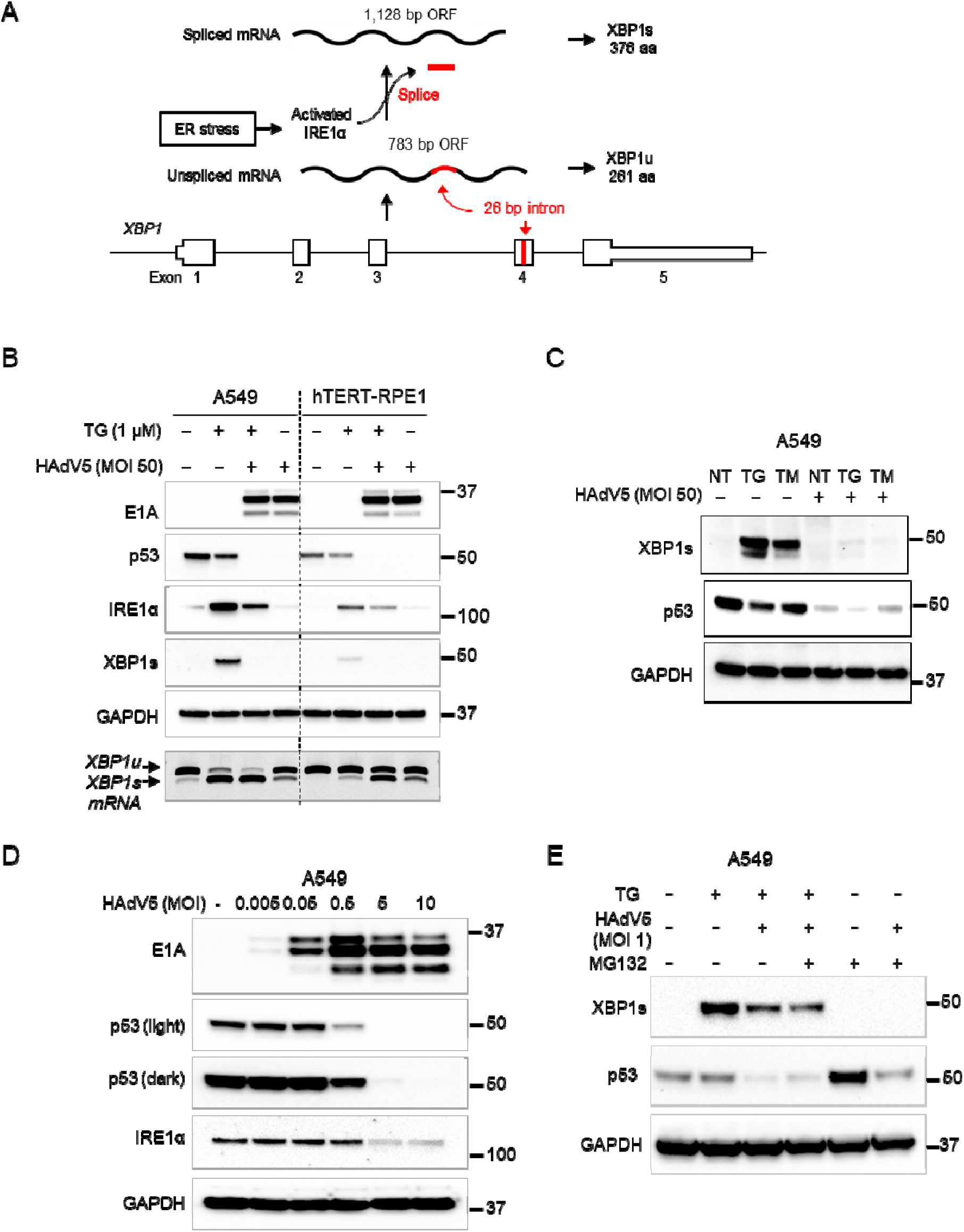
Decreased XBP1s expression after infection with HAdV5. (A) When activated by ER stress, IRE1α excises a 26-bp splice from the *XBP1* transcript, causing a frameshift. As a result, the *XBP1* gene produces two distinct proteins, XBP1u and XBP1s, with unrelated c-termini. (B) A549 and hTERT-RPE1 cells were infected with HAdV5 (MOI=50) for 24 h, in the presence (+) or absence (-) of thapsigargin (TG, 1 µM). The indicated proteins were assessed by immunoblot; the migration of molecular weight markers (kDa) are shown at right. Spliced and unspliced *XBP1* RNAs were reverse transcribed and amplified by PCR. (C) Uninfected or HAdV5-infected A549 cells were treated with 1 µM TG or 5 µg/ml tunicamycin (TM). (D) Cells were continuously exposed to HAdV5 at the indicated MOI for 24 h prior to harvest. (E) Infected and uninfected cells were treated with TG and/or the proteasome inhibitor MG132 for 24 h.

Viral infections commonly result in profound changes in protein flux and thus constitute a strong stimulus for the UPR. A growing body of evidence suggests that the UPR is not merely a passive response to high levels of viral protein expression, but additionally serves as an antiviral “danger signal” that stimulates pattern recognition and paracrine signaling, which coordinately prevent the further spread of infection.^8,9^ Conversely, many viruses, including adenoviruses, influenza A virus, hepatitis C virus, herpesviruses and flaviviruses such as Dengue virus and Zika virus, encode proteins that interact with UPR regulators and thus actively modulate UPR activity.^10^ Some viruses downregulate or suppress the UPR, suggesting that the UPR can attenuate viral propagation. Cytomegaloviruses, for example, prevent IRE1α-XBP1 pathway activation via expression of the M50/UL50 protein, which binds to IRE1α and targets it for degradation.^11^

Other virus-encoded proteins stimulate UPR activation, suggesting that viruses have evolved diverse strategies for circumventing the antiviral functions of the UPR and in some cases adopting the molecular mechanisms that mediate the UPR in order to facilitate their own propagation. The complexity of the UPR sensor-effector network is accordingly mirrored in the viruses that co-opt it. For example, flaviviruses, which have been intensively studied in this regard, sequentially modulate different branches of the UPR during their replication cycle.^12^

The life cycle of HAdV5 also appears to be tightly meshed with the UPR machinery. Adenoviral propagation can be experimentally enhanced by genetic or chemical induction of the UPR.^13^ This effect specifically involves the IRE1α-XBP1 signaling axis and is specifically robust in cancer cells, which are inherently predisposed to ER stress^14^. Recent evidence suggests that HAdV5 can itself potentiate UPR induction through expression of the viral E3-19K protein, which localizes in the ER lumen and activates IRE1α via protein-protein interactions^15^ XBP1s reportedly binds promoters in the HAdV5 E1 and E4 regions and thus stimulates early viral gene expression. The induction of XBP1s by HAdV5 E3-19K is currently understood to be important for viral persistence.^15^

HAdV5 expresses two early proteins, E1B-55K and E4orf6, that together mediate the degradation of host regulatory proteins that restrict viral propagation.^16^ These viral proteins and several cellular proteins form a ubiquitin ligase complex, which targets host restriction factors for degradation by the proteasome. Cellular targets include components of the MRE11-RAD50-NBS1 (MRN) complex ^17^ and the tumor suppressor p53,^18^ which respectively sense the viral double strand DNA genome and prevent its replication during S phase.

In addition to its expansive roles in the regulation of cell growth, cell death and metabolism,^19^ p53 can negatively regulate the UPR. A ternary complex composed of p53, IRE1α and the ER-resident ubiquitin ligase synoviolin (SYVN1) targets IRE1α for proteasomal degradation.^20^ The levels of IRE1α protein and IRE1α-XBP1 signaling are accordingly elevated in p53-deficient cancer cells that are unable to form this degradative complex. Namba *et al.* ^20^ proposed that the attendant increase in IRE1α-XBP1 activity that accompanies the loss of p53 during tumorigenesis allows evolving cell populations to expand their protein folding capacity and thereby enhance their survival.

Based on these recent findings and the events known to occur during the early stages of HAdV5 infection, we hypothesized that the degradation of p53 by E1B-55K/E4orf6 might contribute to the sustained increase in IRE1α-XBP1 activity required for viral persistence. Here, we report that p53 did in fact downregulate IRE1α-XBP1 pathway activation in HAdV5-infected immortalized human cells. However, HAdV5 inhibited the expression of IRE1α and its downstream effector XBP1s following their activation by ER stress. We discuss how these apparently paradoxical effects on the IRE1α-XBP1 signaling axis might coordinately optimize viral replication during the rapid changes in the intracellular environment that accompany viral infection.

## Results

Previous studies have demonstrated that chemically-induced ER stress can promote adenovirus propagation.^13^ To investigate the status of the IRE1α-XBP1 and p53 signaling pathways under these enhanced stress conditions, we first treated uninfected and infected cells with thapsigargin, a small molecule inhibitor of the sarcoplasmic/endoplasmic reticulum calcium ATPase (SERCA). SERCA is a membrane-associated protein complex that transports calcium ions into intracellular organelles; its inhibition leads to the rapid depletion of intralumenal calcium stores in the ER. At a high multiplicity of infection (MOI), HAdV5 robustly expressed E1A protein in both A549, a lung cancer cell line that is highly permissive to HAdV5 infection and hTERT-RPE1 (Fig 1B), an immortalized cell line derived from retinal pigment epithelia that is frequently used to model normal p53 functions.^21^ A549 and hTERT-RPE1 each harbor wild type p53 and exhibit DNA damage-inducible expression of p53 target genes.

IRE1α and XBP1s proteins were increased in both cell lines upon treatment with thapsigargin alone (Fig 1B), as expected. However, HAdV5 infection in the presence of thapsigargin led to a marked decrease in the induction of IRE1α and XBP1s. In contrast, the combination of thapsigargin and HAdV5 had an additive effect on IRE1α-mediated *XBP1* mRNA splicing (Fig 1B, lower panel). To determine whether the observed effects of HAdV5 on IRE1α and XBP1s induction were a general effect of UPR activation rather than a specific interaction with thapsigargin and/or SERCA, we repeated this experiment with tunicamycin, which blocks the first step of glycoprotein synthesis and thereby causes an overabundance of unfolded proteins in the ER. We observed essentially the same result (Fig 1C).

Upon titration of HAdV5 onto A549 cells, p53 was degraded at a lower MOI than what was required for loss of IRE1α expression (Fig 1D). XBP1s was not detectable under these conditions (not shown). The proteasome inhibitor MG132 increased the abundance of p53 in uninfected cells and restored p53 expression after HAdV5 infection, as expected, but had no apparent effect on XBP1s (Fig 1E).

To determine whether the downregulation of XBP1s was dependent on viral gene expression, we employed two recombinant viruses. In one virus (ΔE1-GFP) the entire E1 gene cluster, which is required for viral gene expression and replication, was replaced with a CMV-GFP expression cassette. In the ΔE3-GFP virus, the same GFP cassette was inserted into a deletion in the E3 region, leaving the E1 region intact. As expected, the replication-competent ΔE3-GFP virus expressed increasing amounts of the viral DSB protein, which is required for DNA replication, across a 100-fold range of MOI. This virus eliminated XBP1s expression, while the replication-incompetent ΔE1-GFP virus expressed no detectable DBP and had a minimal effect on XBP1s induction. Notably, the replication-competent virus consistently expressed roughly 10-fold more GFP protein. We confirmed that the titers of the two viruses were similar by immunohistochemical staining with anti-hexon antibody (Fig 2B).

**Figure 2.**
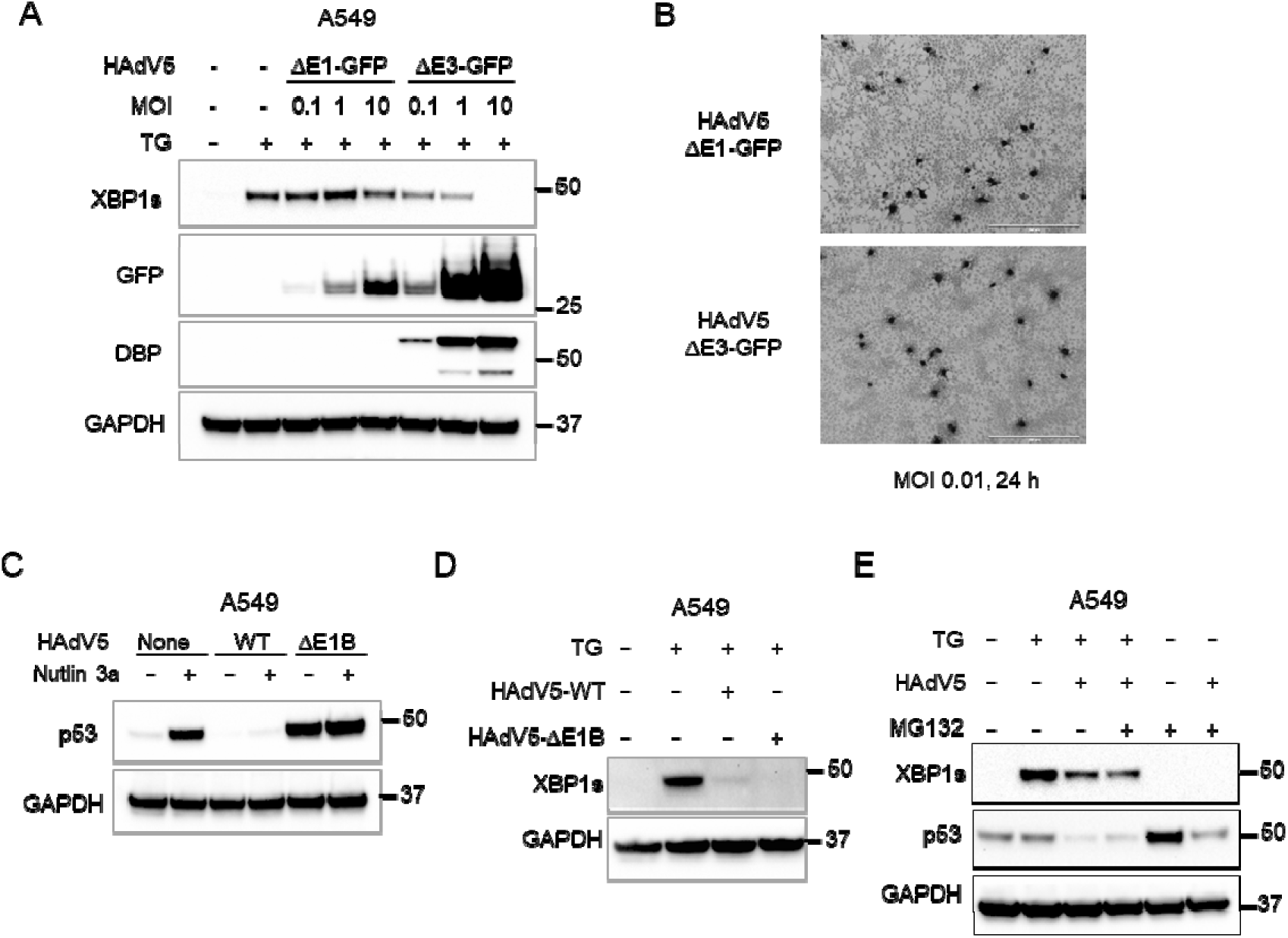
Requirements for XBP1s suppression. (A) A549 cells were incubated with the replication-incompetent HAdV5 virus ΔE1-GFP or the replication-competent virus ΔE3-GFP at the indicated MOI for 24 h. Proteins were harvested for immunoblot analysis. (B) The functional titers of ΔE1-GFP and ΔE3-GFP viruses were determined by immunocytochemistry with an anti-hexon antibody. (C) A549 cells were infected with unmodified HAdV5 (WT) or the *dl*1520 mutan (ΔE1B) that does not express E1B-55K for 24 h (MOI=1), in the presence or absence of the MDM2 inhibitor nutlin-3a (10 µM). Cells were incubated with the indicated viruses (MOI=1): (D) in the presence or absence of 1 µM TG or (E) in the presence or absence of the proteasome inhibitor MG132 (20 µM).

HAdV5 infection activates the DNA damage signaling pathways that control p53 stabilization^22^ but this cellular response is counteracted in part by the E1B-55K protein. As expected, infection with an E1B-55K-deficient virus triggered the stabilization of p53 (Fig 2C). This mutant virus had the same suppressive effect on XBP1s expression as wild type HAdV5 (Fig 2D), providing further evidence that the viral strategies for suppression of p53 and XBP1s proteins are distinct. The level of p53 increased when cells were treated with MG132 alone, consistent with the known mechanism of p53 turnover. MG132 treatment only partially restored p53 expression in the presence of virus, but did not affect XBP1s abundance under any of the conditions tested (Fig 2E).

Spliced viral *E1A* transcripts were readily detectable in infected cells (Fig 3A). *E1A* expression was several-fold higher in A549 compared with hTERT-RPE1 at the same MOI. In A549, treatment with thapsigargin increased the level of E1A expression, a finding that was consistent with prior observations in cancer cells.^13^ *ERN1* encodes the IRE1α protein, an upstream effector as well as a downstream target of the UPR. *ERN1* transcription was strongly induced by thapsigargin. In A549, *ERN1* transcription was lower after HAdV5 infection, but this effect was not apparent in hTERT-RPE1 (Fig 3A).

**Figure 3.**
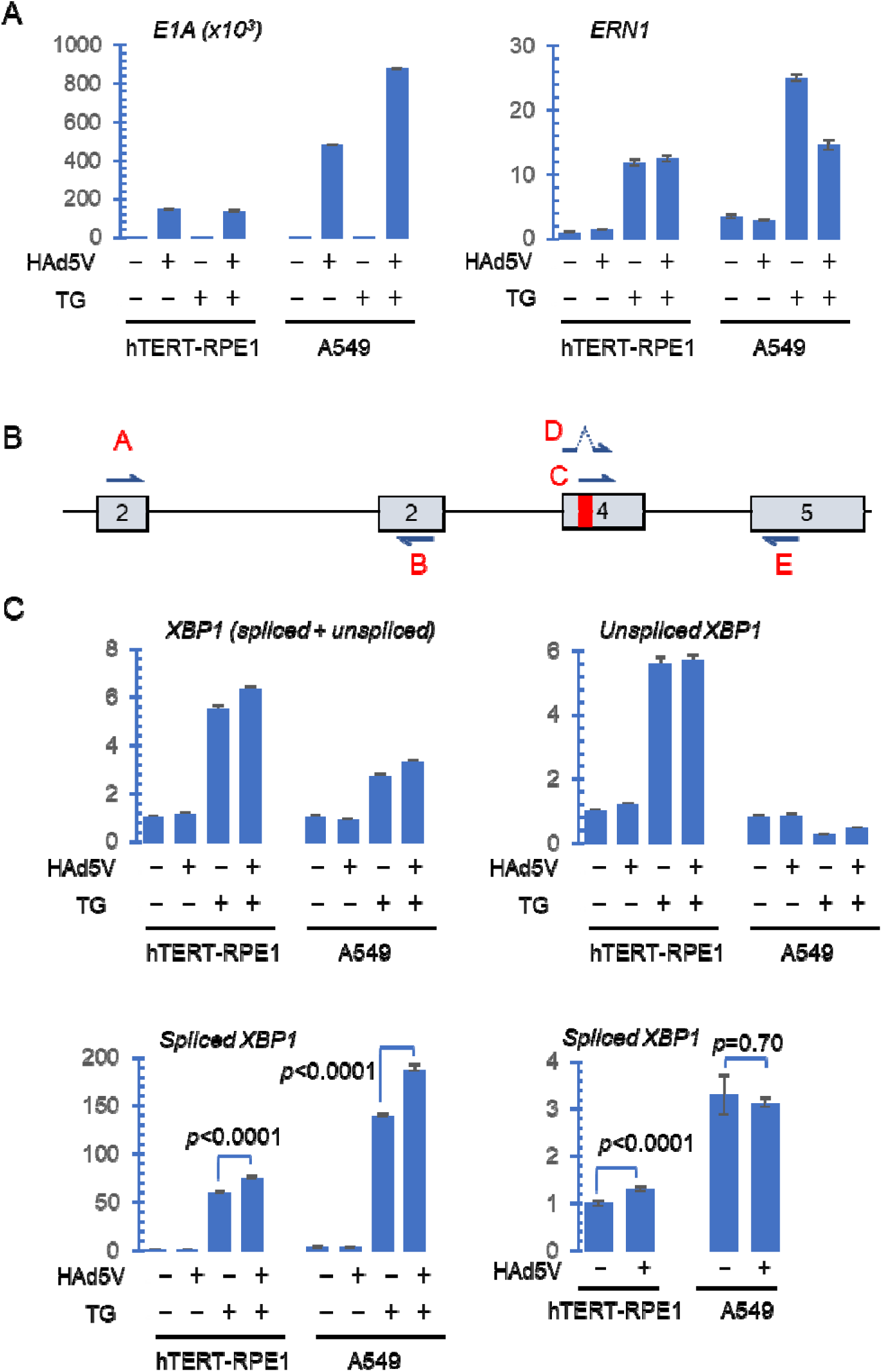
Gene expression and XBP1 RNA splicing after HAdV5 infection. (A) Cells grown in subconfluent monolayers were incubated with wild type HAdV5 (MOI=50) in the presence or absence of 1 µM TG for 16 h. The abundance of viral *E1A* and *ERN1* transcripts were quantified by RT-qPCR. (B) An illustration of *XBP1* exons 2-5, with the excised 26-bp intron shown in red. The primers designated A and B were used to quantify all XBP1 transcripts; the pairs C/E and D/E were used to detect unspliced and spliced XBP1 transcripts, respectively. (D) The abundance of XBP1 transcripts was assessed by RT-qPCR. Error bars represent the standard error of the mean for three replicates. Statistical significance between different samples was determined by unpaired *t*-test, as indicated.

To assess whether the XBP1s protein was downregulated at the transcriptional level after viral infection, we assessed spliced and unspliced *XBP1* mRNAs levels with transcript-specific primers, as previously described^23^ (Fig 3B). Like *ERN1*, *XBP1* is itself a downstream target of the UPR, primarily controlled by the ATF6 branch.^24^ Accordingly, the overall abundance of *XBP1* transcripts was increased by thapsigargin treatment in both cell lines (Fig 3C). The levels of unspliced *XBP1* RNA were increased >5-fold in hTERT-RPE1 after thapsigargin treatment, irrespective of HAdV5 infection, but were little changed in A549. One interpretation of this marked increase in unspliced XBP1 transcripts in the context of UPR activation is that IRE1α activity is limiting in hTERT-RPE1. A549 cells, like many cancer cell types, appear to have a high capacity for *XBP1* splicing.

As expected, spliced *XBP1* transcripts were strongly induced in both cell lines after thapsigargin treatment (Fig 3C). Consistent with our gel analysis of *XBP1* RT-PCR products (Fig 1B) infection of thapsigargin-treated cells led to increased levels of *XBP1* splicing. The abundance of spliced *XBP1* RNAs was slightly increased by HAdV5 infection alone in hTERT-RPE1 but not in A549. This small increase in splicing was consistent with what we observed upon gel electrophoresis of RT-PCR products (Fig 1B).

We used isogenic p53-knockout cells to further investigate how the degradation of p53 by HAdV5 influenced the induction of the UPR and replication of the virus. In both cell lines tested, p53 was degraded after 24 h at a low MOI (Fig 4A, left panels). At this level of virus, p53-deficient hTERT-RPE1 cells exhibited higher levels of E1A expression, increased induction of IRE1α and increased XBP1 splicing compared with p53-proficient cells. In contrast, infection of A549 cells with constitutively upregulated IRE1α caused the downregulation of this protein. Despite this suppressive effect on IRE1α expression, HAdV5 infection nonetheless caused a modest increase in *XBP1* splicing (Fig 4A, right panels). This result is consistent with the increased capacity for *XBP1* RNA splicing in A549, inferred from our transcript analysis (Fig 3D).

**Figure 4.**
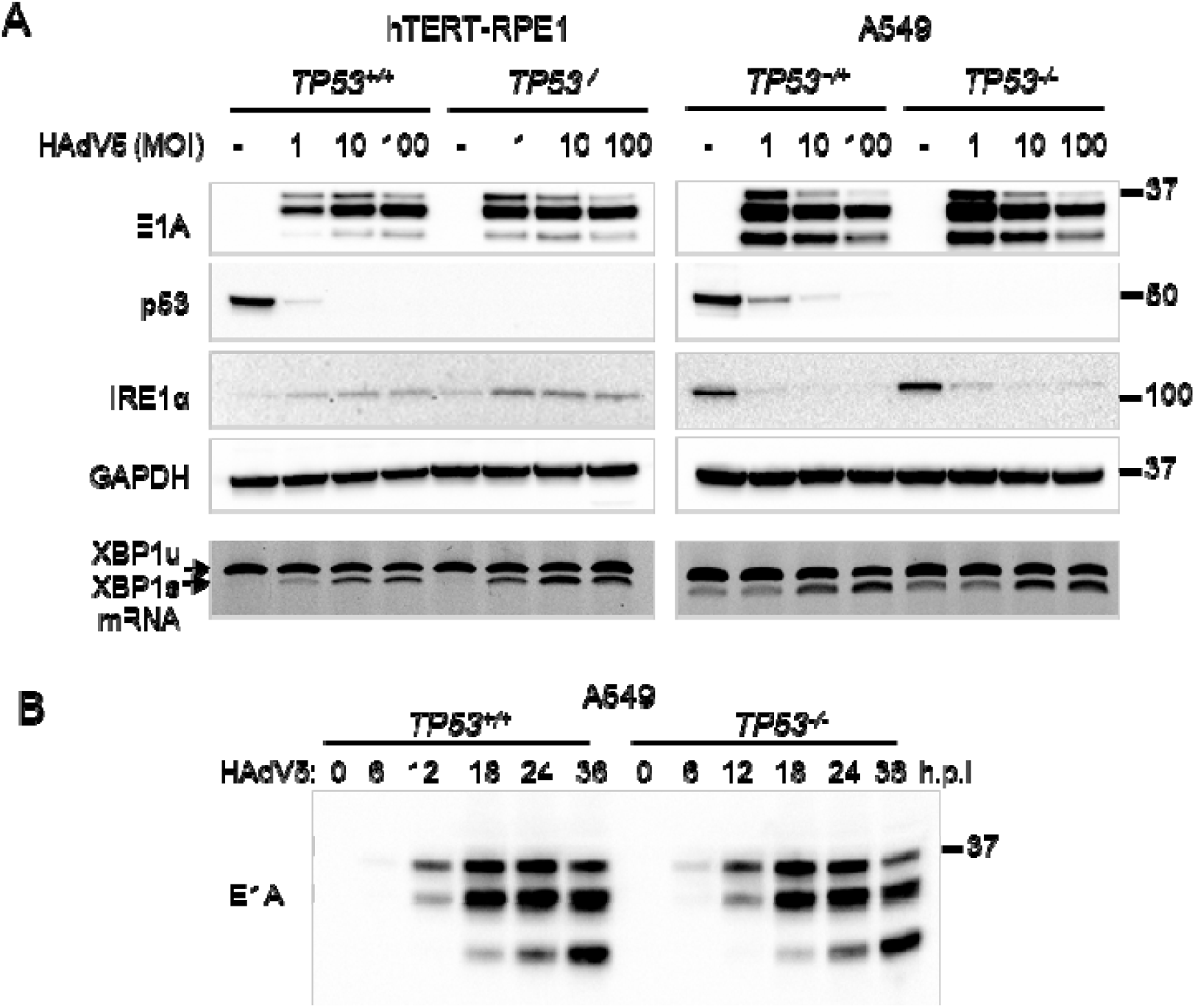
HAdV5 E1A and IRE1α expression in isogenic p53-proficient and -deficient cells. (A) hTERT-RPE1 cells, A549 cells and their corresponding p53-knockouts were incubated with wild type HAdV5 at the indicated MOI. Proteins were assessed by immunoblot. Unspliced (*XBP1u*) and spliced (*XBP1s*) RNAs were extracted from cells treated in parallel and assessed by RT-PCR. (B) Isogenic A549 cells were incubated with wild type HAdV5 (MOI=1) for 36 h. Cells were harvested for immunoblot at the indicated time points after infection (h.p.i, hours post-infection).

At low MOI, viral E1A proteins were more highly expressed in *TP53^-/-^*hTERT-RPE1 compared with the parental cell line, whereas A549 cells supported robust E1A expression irrespective of *TP53* genotype (Fig 4A). The expression of E1A proteins was also similar in isogenic p53-proficient and -deficient A549 cells over a 36 h time course after infection (Fig 4B), further indicating that p53 had little impact on viral gene expression in these cells.

Infection with the HAdV5 mutant *dl*1520, which does not express the E1B-55K protein, resulted in the stabilization of p53 in both cell lines, as expected (Fig 5A); p53 induction was particularly pronounced in A549 cells. In hTERT-RPE1 cells, the expression of E1A proteins from the E1B-attenuated virus was notably lower than from wild type HAdV5 in hTERT-RPE1, whereas A549 supported similarly high levels of protein expression from both viruses. Irrespective of their impact on p53 levels or ability to express E1A, both viruses modestly suppressed the expression of synoviolin, a negative regulator of IREα stability (Fig 5A).

**Figure 5.**
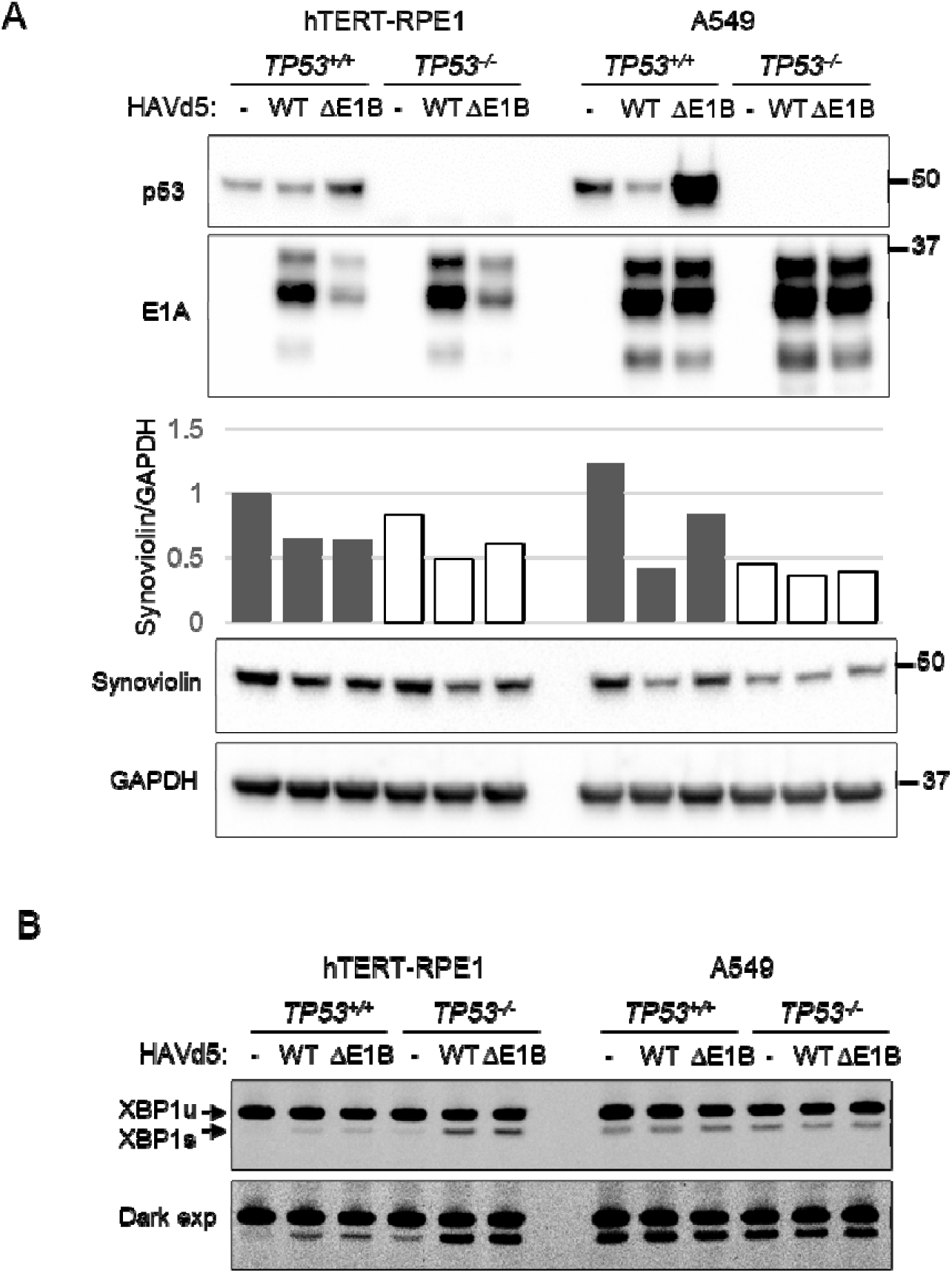
Viral gene expression and *XBP1* RNA splicing after HAdV5 infection. (A) p53-proficient and -deficient cells were incubated with HAdV5 (MOI=1) for 24 h. Protein lysates were probed with the indicated antibodies. The signals for GAPDH and Synoviolin were scanned with ImageJ and plotted as a Synoviolin/GAPDH ratio. (B) Cells treated in parallel were harvested for RNA extraction and RT-PCR analysis. The migration of unspliced (XBP1u) and spliced (XBP1s) cDNAs is indicated by arrows. The lower panel is a darker exposure of the image shown in the upper panel.

Consistent with the decrease in synoviolin, IRE1α activity was induced after HAdV5 infection in hTERT-RPE1 (Fig 4A). Moreover, the interplay between HAdV5, synoviolin and IRE1α is consistent with prior observations of IRE1α induction by the HAdV5 ER-localized E3-19K protein. In p53-deficient hTERT-RPE1 cells, wild type and ΔE1B-55K viruses stimulated *XBP1* mRNA splicing by IRE1α to a similar extent (Fig 5B). In A549 cells, this low MOI did not increase the level of *XBP1* splicing over background, irrespective of *TP53* status.

## Discussion

Cancer and infections are similarly characterized by abnormalities in protein expression. Accordingly, the IRE1α-XBP1 branch of the UPR has been broadly implicated in pathogenesis. A deeper understanding of the UPR may yield new strategies for treating a wide range of common human diseases. In cancers, the upregulation of IRE1α-XBP1 signaling has been proposed as a mechanism for relieving the ER stress associated with oncogene activation.^24^ In the infected cell, the IRE1α-XBP1 pathway appears to be pivotal as both the host and pathogen vie for control over cell fate. The results presented here confirm and extend recent findings regarding the roles of this ubiquitous pathway in cancer and adenovirus infection.

In agreement with previous reports, we found that HAdV5 infection could trigger IRE1α-XBP1 activation. However, the marked suppression of XBP1s induction by HAdV5 is seemingly inconsistent with the proposed role for XBP1s in chronic virus infection and latency. A biphasic pattern of XBP1s regulation could reconcile these findings (Fig 6). In this temporal model, the immediate elimination of p53 by the virus after infection would relieve the p53-mediated suppression of IRE1α and thereby allow XBP1s to be transiently expressed. A brief period of XBP1s activity could promote a rapid increase in E1A-mediated viral gene expression which would in turn facilitate viral replication. The subsequent inhibition of XBP1s could hamper antiviral effects of the UPR that might otherwise be predominant at later points during the viral life cycle. It is possible that a dynamic interaction between the adenovirus and its host, as previously observed in flaviviruses, could model a general feature of many viruses. Additional study will be needed to elucidate such phase-specific interactions between virus and host, and to determine if this biphasic model is more broadly applicable.

**Figure 6.**
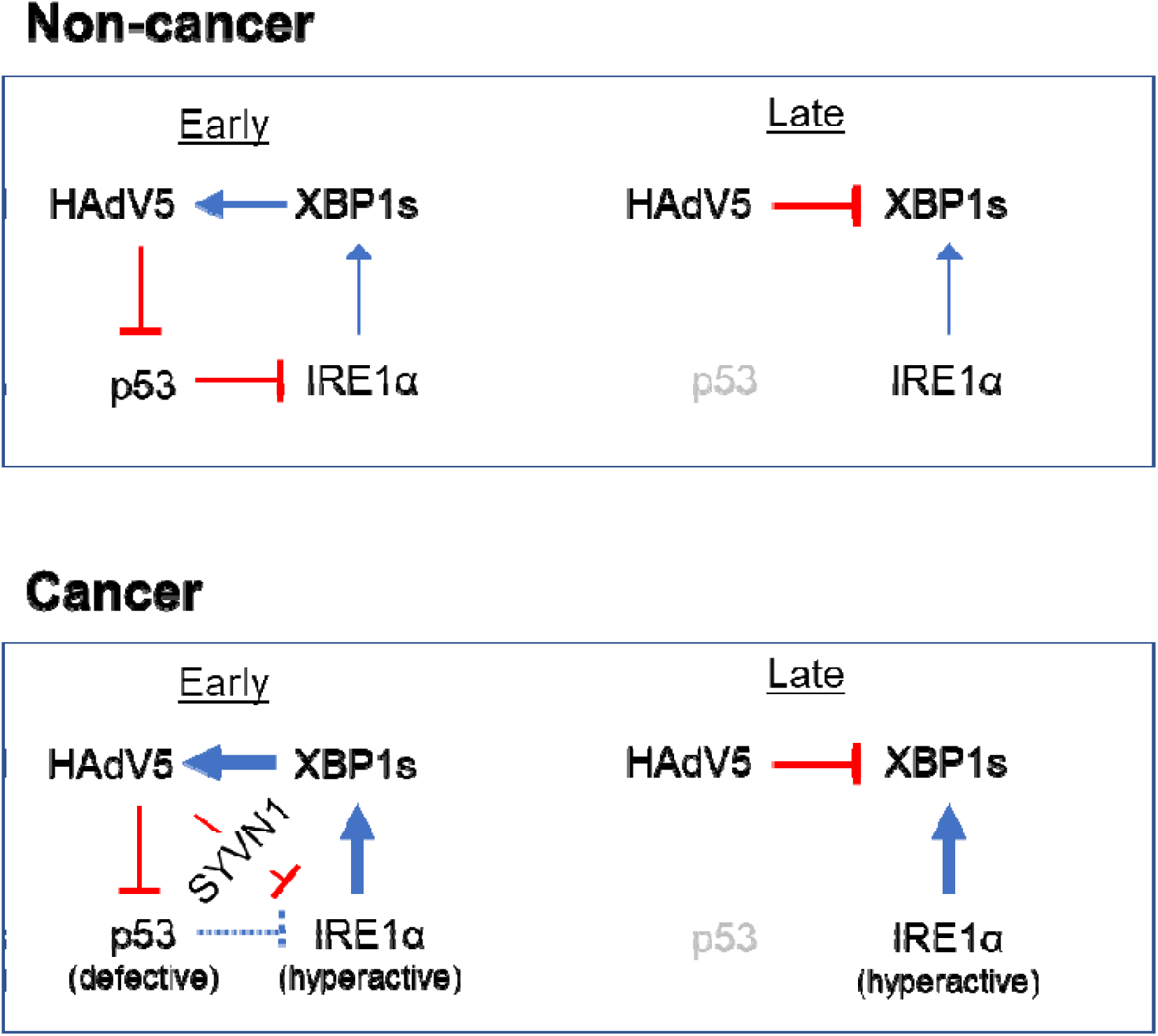
Interplay between HAdV5, p53 and the IRE1α-XBP1 pathway. In this biphasic model, non-cancer cells with intact regulation of the IRE1α-XBP1 pathway first suppress p53 to allow the transient de-repression of IRE1α activity, and subsequently suppress XBP1s translation. Cancer cells constitutively activate IRE1α, either by loss of p53 or another unknown mechanism. Regulation of IRE1α by p53 and other factors may therefore represent an intrinsic barrier to infection that is lost during tumorigenesis. See text for additional details.

Although the precise mechanistic basis for the downregulation of XBP1s after HAdV5 infection remains unclear, our results indicate that the change in XBP1s abundance occurs post-transcriptionally and that the loss of protein expression is dependent neither on E1B-55K nor the proteasome. In the absence of targeted degradation, the loss of XBP1s may be related to the global shutdown of host protein synthesis that is orchestrated by HAdV5.

This study highlights important differences between cancer cells and normal (non-transformed) human epithelial cells. The hTERT-RPE1 cell line was originally derived from primary cells that have been immortalized *in vitro*, and is commonly employed as a surrogate for normal epithelial cell types. When comparing hTERT-RPE1 cells with A549, a highly permissive line frequently employed as an experimental viral host, we found that IRE1α-XBP1 signaling was tightly regulated in the former but relatively deregulated in the latter. This observation is in accordance with the current consensus view that the UPR is frequently abnormal in cancers. For this study, we chose to focus on two cell lines that are widely used for studying p53 and viral infections. In the future, it will be important to determine whether the divergent patterns of IREα-XBP1 signaling in response to viral infection observed here will be generalizable to diverse cell types and model systems.

One important difference between many cancer cells and normal cells is the loss of p53 function. A recent study from our laboratory has provided evidence that strong *in vivo* selection against p53 activity during tumorigenesis inevitably results in functional p53 defects among virtually all cancer cells, even in those that retain wild type *TP53* alleles.^21^ We reported that the hTERT-RPE1 model, in which p53-dependent senescence was bypassed via the forced expression of telomerase, retains a wide variety of robust p53 functions that are often difficult to appreciate in other human cell models. The data presented here suggest that the hTERT-RPE1 system provides a unique view into the dynamic interactions between HAdV5, p53 and the IRE1α-XBP1 signaling pathway. Conversely, the constitutive upregulation of IREα in A549 cells could reflect the circumvention of p53-mediated IREα suppression during the evolution of the original lung tumor.

## Materials and Methods

### Cell lines and cell culture

Parental A549 cells were purchased from ATCC. The puromycin-sensitive hTERT-RPE1 cell line was a gift from Andrew Holland. Cells were routinely grown at 37°C in 5% CO_2_ in DMEM/F12 supplemented with 10% fetal bovine serum (FBS) and penicillin/streptomycin. The parental cell line and all derivatives were authenticated by Short Tandem Repeat profiling and tested for the presence of mycoplasma at the Johns Hopkins Genomic Resources Core Facility.

### Viruses

Unmodified HAdV5 was purchased from ATCC. The *d1*520 mutant was a gift from Patrick Hearing (Stony Brook University). The ΔE1-GFP and ΔE3-GFP viruses were assembled from modular components, as described.^25^ Virus titers were determined with the QuickTiter Adenovirus Immunoassay kit (Cell Biolabs).^20^

### Generation of TP53^-/-^ cells

The procedure for generating p53-knockout derivatives of hTERT-RPE1 cells were recently described.^21^ The same strategy was used to generate homozygous deletions of *TP53* exon 1 in A549. Briefly, the flanking guides 5’-TAGTATCTACGGCACCAGGT-3’ and 5’-TCAGCTCGGGAAAATCGCTG-3’ were designed to create a 385 bp deletion that included the entire exon. Oligonucleotide duplexes were directly cloned into the plasmid vector pSpCas9(BB)-2A-Puro (pX459) V2.0, a gift from Feng Zhang (Addgene #62988). Plasmids were co-transfected with Lipofectamine 3000 (ThermoFisher Scientific). Following a 4 d selection in 2 µg/ml puromycin, the surviving cells were plated to limiting dilution in 96-well plates. Individual subclones were expanded and screened by western blot. Multiple knockout clones were identified and confirmed by PCR across the exon 1 deletion, using the forward primer 5’-CTCCAAAATGATTTCCACCAAT-3’ and the reverse primer 5’-ACTTTGAGTTCGGATGGTCCTA-3’.

### Western blots, antibodies

Non-denatured protein extracts were prepared in Cell Lysis Buffer (Cell Signal Technologies). resolved on Bolt Bis-Tris minigels (ThermoFisher Scientific) and transferred to PVDF membranes (MilliporeSigma). Antibodies for the detection of p53 (DO-1) and HAdV2/5 E1A (M73) were obtained from Santa Cruz Biotechnology. Antibodies recognizing IRE1α (14C10) and XBP1s (E8C2Z) were purchased from Cell Signaling Technology. Polyclonal antibodies for the detection of GAPDH and synoviolin were purchased from MilliporeSigma and Bethyl, respectively. Hybridoma supernatant for the detection of HAdV5 DBP was a gift from David Ornelles (Wake Forest University).

### Drug treatments

The MDM2 inhibitor nutlin-3a and the proteasome inhibitor MG-132 were purchased from Enzo Life Sciences, dissolved in DMSO and used at final concentrations of 10 µM and 20 µM respectively. The SERCA inhibitor thapsigargin was purchased from Cell Signaling Technology and dissolved in DMSO.

### XBP1 splicing assay

Cells growing in monolayer cultures were directly harvested in RNA lysis buffer and processed with the Monarch RNA purification kit (NEB). cDNA was synthesized with the LunaScript RT SuperMix kit (NEB) from approximately 500 ng of total RNA. *XBP1* cDNAs were amplified in a PCR reaction containing 2X Phusion Master Mix (NEB). The forward primer 5’-GGAGTTAAGACAGCGCTTGGGGA-3’ and the reverse primer 5’-TGTTCTGGAGGGGTGACAACTGGG-3’, were previously described.^20^ Reaction products were separated on 6% Tris-Borate-EDTA minigels (ThermoScientific), stained with ethidium bromide and visualized on a GelDoc imaging workstation (BioRad).

### Analysis of gene expression

mRNA expression was assessed by reverse transcription-quantitative PCR (RT-qPCR). Total RNA was extracted from subconfluent cells and treated with DNase I with the Monarch total RNA purification kit (NEB). Reverse transcription and PCR amplification were performed with the Luna One-Step RT-qPCR kit (NEB). Real time PCR amplification was performed on a BioRad CFX96 Real-Time PCR detection system. The primer sets used were as follows: *GAPDH,* forward 5’-GAGTCAACGGATTTGGTCGT-3’, reverse 5’-TTGATTTTGGAGGGATCTCG-3’; *E1A*, forward 5’-CACGGTTGCAGGTCTTGTCATTAT-3’, reverse 5’-GCTCAGACACAGGACTGTA-3’; *ERN1*, forward 5’-CCGAACGTGATCCGCTACTTCT-3’, reverse 5’-CGCAAAGTCCTTCTGCTCCACA-3’; *XBP1* (spliced and unspliced) forward 5’-CTGCCAGAGATCGAAAGAAGGC, reverse 5’-CTCCTGGTTCTCAACTACAAGGC; *XBP1*-spliced, forward 5’-TGCTGAGTCCGCAGCAGGTG-3’, reverse 5’-GCTGGCAGGCTCTGGGGAAG-3’; *XBP1*-unspliced, forward 5’-TGCACCTCTGCAGCAGGTG-3’, reverse 5’-GCTGGCAGGCTCTGGGGAAG-3’. Primers for the specific detection of the spliced and unspliced *XBP1* transcripts were previously described. The abundance of each gene was normalized to the *GAPDH* expression in the same sample.

## Acknowledgements

The authors thank Paul Hwang (NHLBI) for his critical reading of this manuscript. This work was funded by grants from the NIGMS (R01GM135485) and the Foundation for the National Institutes of Health (FNIH).

